# Single-cell RNA sequencing highlights a reduced function of natural killer and cytotoxic T cell in recovered COVID-19 pregnant women

**DOI:** 10.1101/2022.08.18.504053

**Authors:** Nor Haslinda Abd Aziz, Madhuri S. Salker, Aditya Kumar Lankapalli, Mohammed Nasir Shafiee, Ersoy Kocak, Surya Sekhar Pal, Omer Khalid, Norhana Mohd Kasim, Aida Kalok, Norashikin Abdul Fuad, Stephan Ossowski, Nicolas Casadei, Deutsche COVID-19 OMICS Initiative (DeCOI), Sara Y Brucker, Olaf Riess, Yogesh Singh

**Author notes:** Equal contributions (Nor Haslinda Abd Aziz, Madhuri S. Salker). Lead Contact for the study Dr Yogesh Singh, Institute of Medical Genetics and Applied Genomics, NGS Competence Centre Tübingen (NCCT), Research Institute of Women’s Hospital, University of Tübingen, Calwerstraße 7, 72076, Tübingen, Germany.

## Abstract

Pregnancy is a complex phenomenon during which women undergo immense immunological change throughout this period. Having an infection with the SARS-CoV-2 virus leads to an additional burden on the highly stretched immune response. Some studies suggest that age-matched pregnant women are more prone to SARS-CoV-2 infection compared with normal healthy (non-pregnant) women, while alternative evidence proposed that pregnant women are neither susceptible nor develop severe symptoms. This discrepancy in different findings regarding the immune responses of pregnant women infected with SARS-CoV-2 virus is not well understood. In this study, we investigated how SARS-CoV-2 viral infection could modulate the immune landscape during the active infection phase and recovery in pregnant females. Using flow cytometry, we identified that intermediate effector CD8^+^ T cells were increased in pregnant women who had recovered from COVID-19 as opposed to those currently infected. Similarly, an increase in CD4^+^ T helper cells (early or late) during the recovered phase was observed during the recovery phase compared with infected pregnant women or healthy pregnant women, whilst infected pregnant women had a reduced number of late effector CD4^+^ T cells. CD3^+^CD4^-^ CD8^-^NKT cells that diminished during active infection in contrast to healthy pregnant women were significant increase in recovered COVID-19 recovered pregnant women. Further, our single-cell RNA sequencing data revealed that infection of SARS-CoV-2 had changed the gene expression profile of monocytes, CD4^+^ effector cells and antibody producing B cells in convalescent as opposed to healthy pregnant women. Additionally, several genes with cytotoxic function, interferon signalling type I & II, and pro- and anti-inflammatory functions in natural killer cells and CD8^+^ cytotoxic T cells were compromised in recovered patients compared with healthy pregnant women. Overall, our study highlights that SARS-CoV-2 infection deranged the adaptive immune response in pregnant women and could be implicated in pregnancy complications in ongoing pregnancies.

## Introduction

Corona virus disease-19 (COVID-19) is caused by SARS-CoV-2 virus which has led to a global pandemic since its emergence in Wuhan, China in late 2019 (Andersen *et al*, 2020; Zhou *et al*, 2020). More than 6.39 million people have died due to COVID-19 infection and more than 574.9 million people had been infected thus far (until 29.07.2022) (Medicine, 2022). However, these numbers are continuously rising due to newly emergent variants of this virus despite inoculation of COVID-19 vaccine worldwide (Kampf, 2021; Subramanian & Kumar, 2021).

Pregnancy is a complex phenomenon during which women undergo immense immunological changes throughout this period (Abu-Raya *et al*, 2020). During gestation several physiological changes in the anatomical structure of the respiratory system as well as in the immune system could pose a greater risk for the pregnancy complications in women (Bhatia & Chhabra, 2018). The immune system in pregnancy is always in a delicate balance; it must protect the semi-allogenic fetus from maternal rejection whilst simultaneously provide proper fetal development and protection against foreign pathogens (Aghaeepour *et al*, 2017; Watanabe *et al*, 1997). Epidemiological findings from other pandemics (e.g. influenza, Zika and Ebola) highlighted that pregnant women are more susceptible to severe complications and death from viral infection (Alberca *et al*, 2020; Silasi *et al*, 2015; Wilder-Smith, 2021). Having an infection with SARS-CoV-2 virus could hamper the over stretched immune response. Though, the real time number of infections in pregnant women appeared to be neglected due to limited studies (Nidhi, 2021). Global maternal and fetal outcomes have worsened during the COVID-19 pandemic such as increase in maternal death, pre-term labour, stillbirth, ruptured ectopic pregnancies and maternal depression (Henarejos-Castillo *et al*, 2020; Liu *et al*, 2020; Villar *et al*, 2021). Recent studies suggested that age matched pregnant women are more prone to SARS-CoV-2 infection compared with healthy non-pregnant women (Villar *et al*., 2021). SARS-CoV-2 infection in pregnant mothers lead to increased placental inflammation, predisposition to the development of maternal vascular thrombosis, higher caesarean rate, fetal growth restriction and increased risk of preterm delivery (Wong *et al*, 2022). Additionally, altered villous maturation and severe-critical maternal COVID-19 infection were associated with an elevated risk of poor Apgar scores at birth and maternal mortality, respectively (Wong *et al*., 2022). In contrast, other findings proposed that pregnant women are mostly asymptomatic (90%) and develop mild symptoms after SARS-CoV-2 infection (Mullins *et al*, 2020; Waghmare *et al*, 2021). However, the comparisons are based on small cohort and no global data is available on how the immune response of infected and recovered women is shaped by SARS-CoV-2 virus. A handful of studies investigated how the virus modules the function of immune cells during pregnancy (Bordt *et al*, 2021; Chen *et al*, 2021a; Chen *et al*, 2021b; Lu-Culligan *et al*, 2021; Ovies *et al*, 2021). Thus, there is no clear understanding why there is such a discrepancy in the immune response of pregnant women infected with SARS-CoV-2 virus in different published studies. In our study, we thus applied single-cell RNA sequencing (scRNA-seq) and flowcytometry to investigate how SARS-CoV-2 viral infection could modulates the immune response during the active SARS-CoV-2 infection and after the recovery from COVID-19 disease in pregnant women.

## Results

### Demographics of the pregnant women

A total of 19 pregnant Malaysian and Malaysian Indian pregnant women were enrolled for this study comprised of pregnant healthy controls (Preg-HC; n=11), pregnant SARS-CoV-2 infected (Preg-SARS-CoV-2; n=4), and the recovered from the SARS-CoV-2 infection (Preg-R; n=4), (Table 1) some of the same patients described in our recent published study (Cao *et al*, 2022). In Cao et al, 2022, we used plasma and PBMCs for intracellular cytokine staining and scRNA-seq and immunophenotyping is part of this study. Most of the infected pregnant women were either asymptomatic or some had mild/moderate manifestations of COVID-19. No severe form of COVID-19 was reported and used in this study. Most of the study participants were in third trimester including healthy control pregnant (except one participant which was in the second trimester), infected pregnant (except one participant which was in the second trimester) and recovered pregnant women (except one participant was in the second trimester) (Table 1).

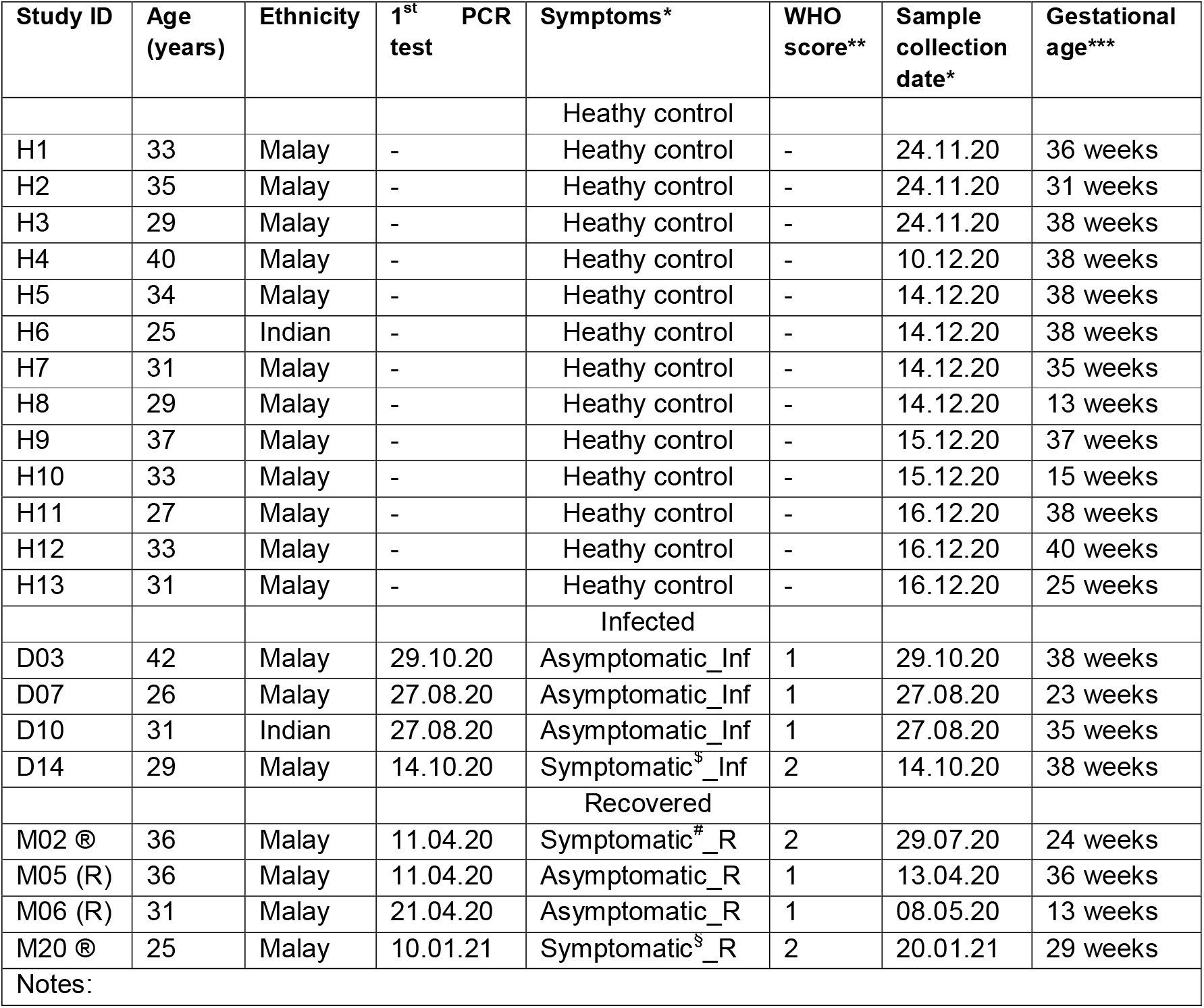
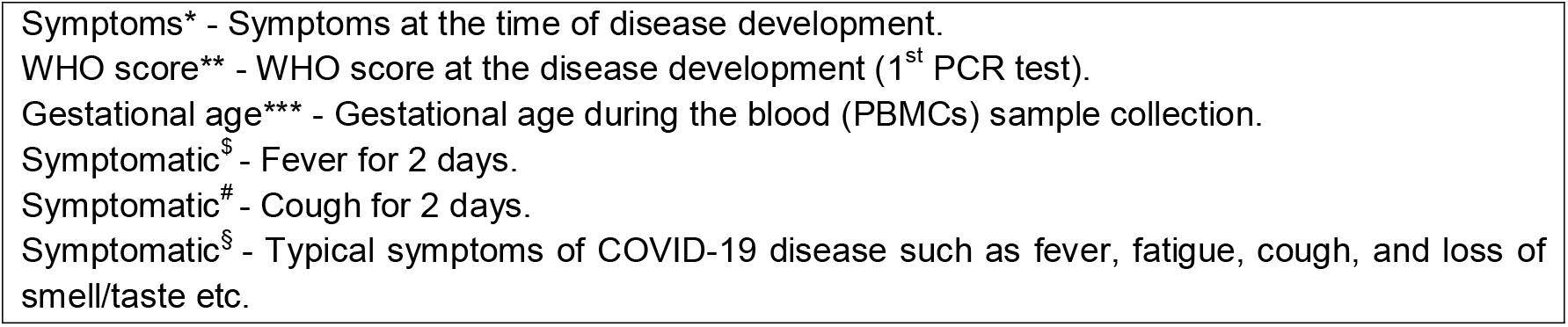

### Immunophenotyping of pregnant, pregnant infected and pregnant recovered women

In our cohort study most of the pregnant women were either infected or recovered and were asymptomatic or mild/moderately affected which is consistent with other previously reported findings (Waghmare *et al*., 2021). We first explored the percentage of different immune subsets in peripheral blood mononuclear cells (PBMCs) to understand the distribution of immune cells in pregnant women. To understand this, we used 14-colour antibody panel to identify the different subtypes of T, B, NK cells and monocytes using flow cytometry. Gating strategy of live lymphocytes and monocytes is displayed in Supp. Fig.1. Overall, 200,000 cells were acquired for each sample and live cells were used for further analysis, we found significantly reduced lymphocytes (p=0.02 Preg-R vs Preg-HC; Kruskal-Wallis nonparametric test (p=0.01) and multiple comparisons based on post-hoc Dunn’s test) in Preg-R patients compared with Preg-HC. Further, a reduced tendency in monocytes in Preg-SARS-CoV-2 and Preg-R patients compared with Preg-HC (Suppl. Fig. 1a, b). No obvious difference was observed either in CD4^+^ or CD8^+^ T cells among Preg-SARS-CoV-2, Preg-R patient samples, and Preg-HC, although, there was a tendency of decreased CD8^+^ T cells and increased CD4^+^ T cells in Preg-SARS-CoV-2 compared with Preg-R or Preg-HC, respectively (Suppl. Fig. 1c).

#### Early and late effector CD4^+^ or CD8^+^ T cells were dysregulated in Preg-SARS-CoV-2 and Preg-R

Further, all the samples were concatenated, and subjected to unsupervised clustering analysis using uniform manifold projection and approximation (UMAP) to classify the clustering of immune cells and identify the difference in immune cell subsets (Fig. 1a, b). We observed 5 major clusters of cells based on 14-colour flow parameters which includes monocytes, CD4^+^ T cells, CD8^+^ T cells, CD19^+^ B cells and CD56^+^ NK cells subsets. Further, using supervised clustering of CD8^+^ T cells, we identified 5 different subsets of CD8^+^ T cells (Fig. 1e). We found that naïve CD8^+^ T cells were reduced in Preg-R patients compared with Preg-HC group and was almost significant (p=0.07) (Fig. 1f). Intermediate effector CD8^+^ T cells were also significantly reduced in Preg-SARS-CoV-2 compared with Preg-R (p=0.03). Late effector CD8^+^ T cells were also significantly reduced in Preg-SARS-CoV-2 compared with Preg-R (p=0.08), however, it did not reach to a significant level (Fig. 1f). This apparent effect was also observed in cell distribution UMAP analysis of CD8^+^ T cells in Preg-SARS-CoV-2 compared with Preg-R (Fig. 1). Further, CD4^+^ T cells were extracted and subject to unsupervised UMAP clustering analysis. We observed that naïve T cells were reduced in Preg-SARS-CoV-2 and Preg-R compared with Preg-HC (Fig. 2a). Based on CD45RA and CCR7 markers, we found that early effector cells tended to increase in Preg-R compared with Preg-HC (p=0.07), though, it did not reach to significant level (Fig. 2b), however, late effector cells were increased in Preg-R compared with Preg-HC (Fig. 2b). Thus, it appears that naïve cells were reduced, whilst intermediate or later effector memory cells were increased in Preg-SARS-CoV-2 and Preg-R compared with HC.

**Fig.1.**
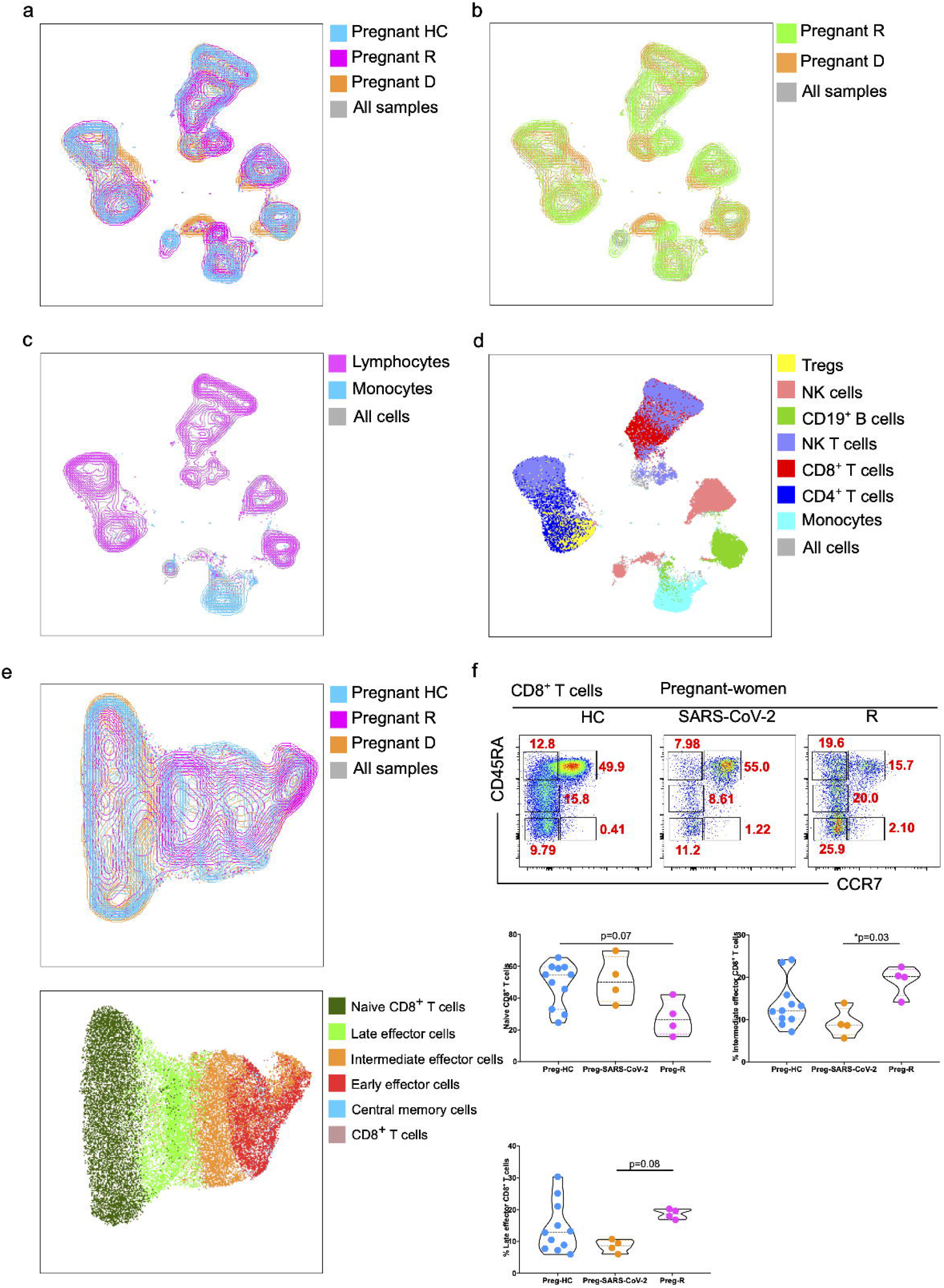
Immunophenotyping of PBMCs in pregnant SARS-CoV-2 infected and recovered patients a. Unsupervised clustering of immune cells based on 14-colour flow cytometry panel. Two major clusters for lymphocytes and monocytes population using UMAP dimensional reduction method. b. Supervised clustering of PBMCs based on gating strategy identified 7 main subsets of cells as shown in a colour coded UMAP plot including monocytes, CD3^+^CD4^+^ T cells, CD3^+^CD8a^+^ T cells, CD3^+^CD56^+^NKT cells CD3^-^CD19^+^ B cells, CD3^-^CD56^+^NK cells and FOXP3^+^ Tregs. c. Overlay of UMAP to show the Preg-HC (Cyan), Preg-R (pink) and Preg-D (orange). All the combined cells were shown in the background (grey). d. Comparisons of immune cells in recovered and infected pregnant women. e. Dynamics of cytotoxic CD4^+^ T cells. Identification of naïve, memory and effector memory CD4^+^ T cells based on CD45RA and CCR7 markers. FACS plots show the CD45RA^+^CCR7^+^ naïve, CD45RA^-^CCR7^+^ CM cells, CD45RA^+high^CCR7^-^ late EM, CD45RA^+mid^CCR7^+^ intermediate EM, CD45RA^-^ CCR7^-^ early EM cells (upper FACS plots). The percentage of naïve, intermediate, and late EM cells shown in violin plots (lower panels). P values show the significance among Preg-HC, Preg-SARS-CoV-2 and Preg-R groups and compared using Wilcoxon test. P value <0.05 considered significant. f. UMAP analysis of total CD4^+^ T cells. UMAP overlay show the Preg-HC (Cyan), Preg-R (pink) and Preg-D (orange). g. UMAP analysis for different CD4^+^ T helper cells based on supervised clustering.

**Fig. 2.**
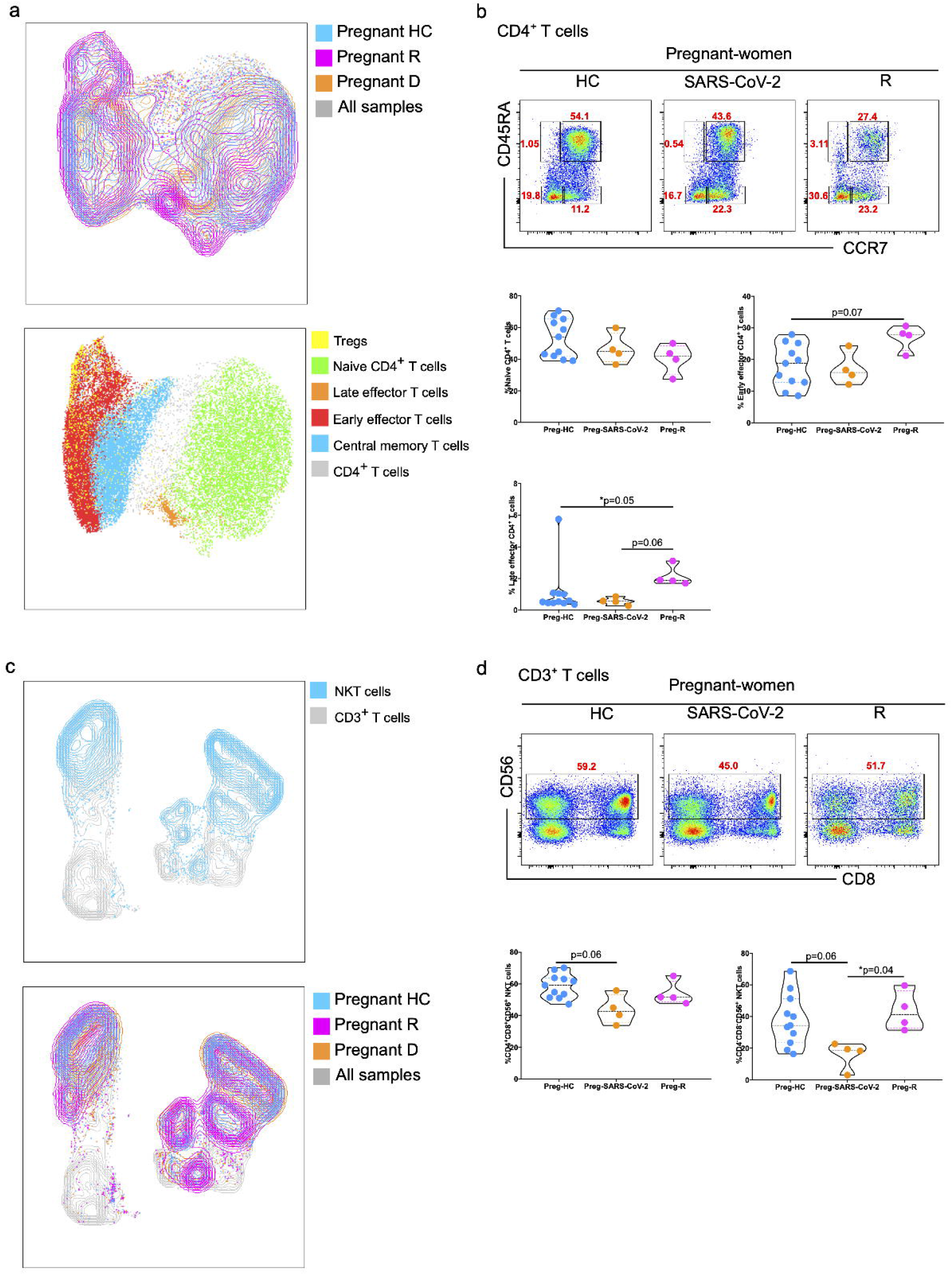
Dysregulated cytotoxic CD8^+^ T cells and NKT cells in infected and recovered pregnant women a. Dynamics of cytotoxic CD8^+^ T cells. Identification of naïve, memory and effector memory cytotoxic CD8^+^ T cells based on CD45RA and CCR7 markers. FACS plots show the CD45RA^+^CCR7^+^ naïve, CD45RA^-^CCR7^+^ CM cells, CD45RA^+high^CCR7^-^ late EM, CD45RA^+mid^CCR7^+^ intermediate EM, CD45RA^-^CCR7^-^ early EM cells (upper FACS plots). The percentage of naïve, intermediate, and late EM cells shown in violin plots (lower panels). P values show the significance among Preg-HC, Preg-SARS-CoV-2 and Preg-R groups and compared using Wilcoxon test. P value <0.05 considered significant. b. UMAP analysis of total CD8a^+^ T cells. UMAP overlay show the Preg-HC (Cyan), Preg-R (pink) and Preg-D (orange). c. UMAP analysis for different cytotoxic T cells based on supervised clustering. d. UMAP plots show the distribution of NKT cells in CD4^+^ and CD8a^+^ T cell compartments. e. FACS plots represent the expression of CD56 on CD3^+^CD8^+^ T cells (Upper FACS panel). The percentage of NKT cells displayed by violin plots among Preg-HC, Preg-SARS-CoV-2 and Preg-R groups and compared using Wilcoxon test. P value <0.05 considered significant. f. Overlay of UMAP to show the Preg-HC (Cyan), Preg-R (pink) and Preg-D (orange) for NKT cells. All the combined samples were shown in background (grey).

#### Decreased CD3^+^CD4^-^CD8^-^ NKT and NK cells in Preg-SARS-CoV-2 or Preg-R

Furthermore, the analysis of PBMCs showed reduced frequency of CD3^+^CD56^+^ NKT cells in Preg-SARS-CoV-2 infected women compared with Preg-HC (Fig. 2d, bottom, left). Interestingly, CD3^+^CD4^-^CD8^-^CD56^+^ NKT cells were significantly reduced in Preg-SARS-CoV-2 infected women compared with Preg-R (Fig. 2d, bottom, right). UMAP analysis depicted a clear distinction of NKT cells in Preg-SARS-CoV-2, Preg-R and Preg-HC. Further CD3^-^CD56^+^ NK cells were characterised using 2D FACS plots and unsupervised UMAP analysis. We observed that CD3^-^CD19^-^CD56^+^HLA-DR^-^ NK cells were reduced in Preg-SARS-Co or Preg-R compared with HC (Fig. 3a, left). However, HLA-DR^+^CD56^+^ NK cells tended to be higher in Preg-SARS-CoV-2 or Preg-R compared with HC (Fig. 3a, right) and similarly, unsupervised clustering of CD3^-^CD19^-^ cells yielded a similar trend (Fig. 3b). Finally, we gated the CD3^-^CD19^-^ CD56^+^HLA-DR^-^ NK and divided the NK cells in different subset using CD16 makers and found that mature CD56^+^CD16^+^ NK cells were decreased in Preg-SARS-CoV-2 or Preg-R compared with HC (Fig. 3c), the difference seen was not significant. Furthermore, non-NK cells were increased in Preg-SARS-CoV-2 and Preg-R (Fig. 3b, c). Overall, our data revealed that mature NK cell numbers were reduced, whilst increased number of activated NK cells in Preg-SARS-CoV-2 and Preg-R compared with Preg-HC.

**Fig. 3.**
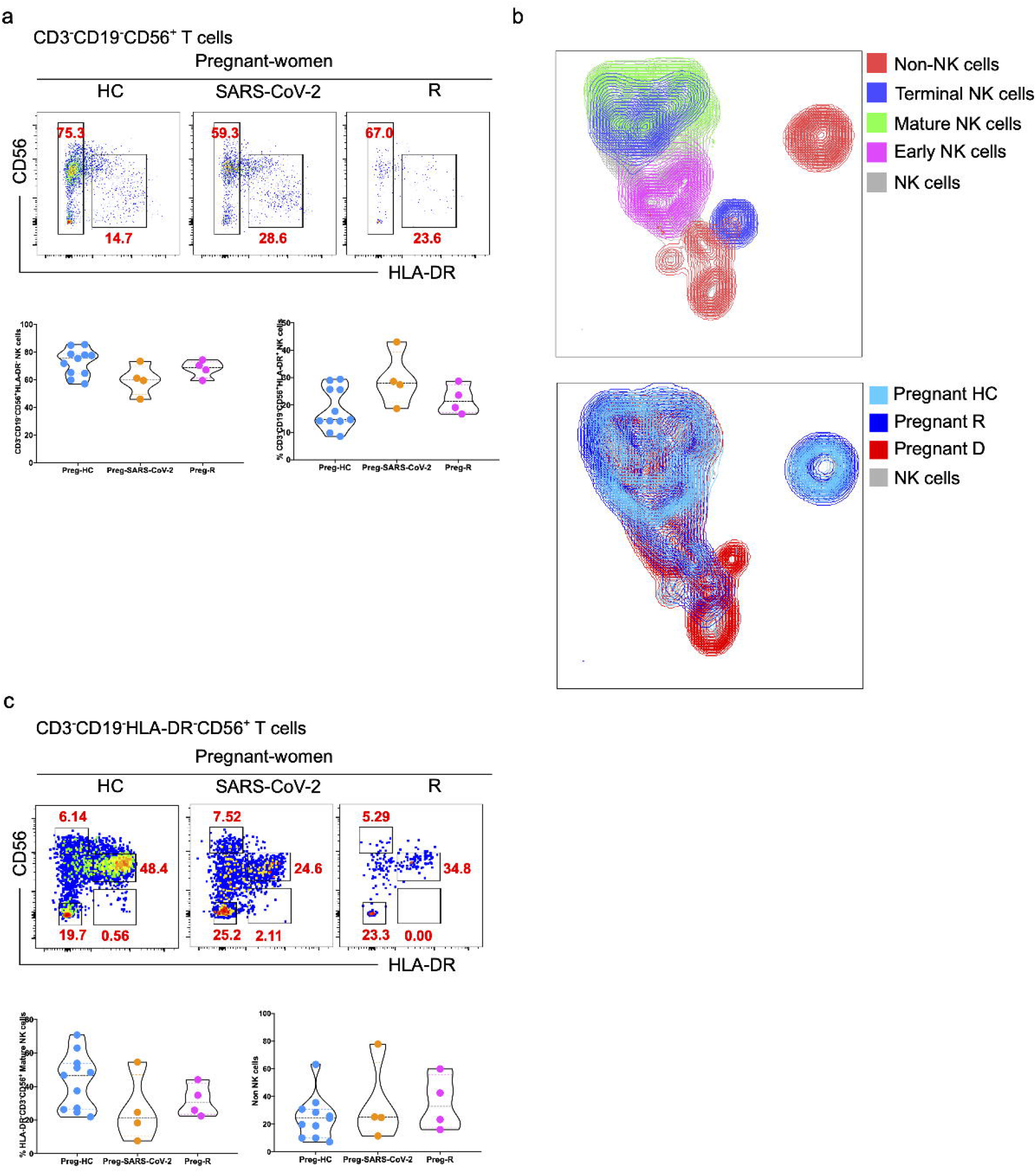
Decreased NK cells in infected and recovered pregnant women a. The percentage of CD3^-^CD19^-^CD56^+^HLA-DR^-^ NK cells shown in FACS plots (upper panel) and violin plots (lower panel) in Preg-HC, Preg-SARS-CoV-2 infected and Preg-R samples. b. UMAP analysis of CD3^-^CD19^-^ cells for different subsets of NK cells based on CD56 and CD16 expression for the terminal, mature, early NK and non-NK cells, while grey coloured cells show the all the samples containing NK cells (upper panel). UMAP overlay for different comparative groups (lower panel). c. The percentage of different subsets of NK cells based on CD56 and CD16 makers. Cells were gated on CD3^-^CD19^-^CD56^+^HLA-DR^-^ for data acquisition. Upper panels show the expression of CD56 and CD16 and lower panel show the violin plots of different NK subsets including mature, intermediate, and terminal NK cells.

### Single cell gene expression profiling of Preg-R and Pre-HC

Flow cytometry results gave an insight at protein level, nonetheless, due to limitation of fluorochromes it is not possible to understand the expression of other molecules which might be affected in the Preg-R patients. We, therefore, employed single cell RNA-sequencing (scRNA-seq) gene expression profiling using the 10x chromium Gel Bead-in-Emulsion (GEMs) to verify this. We used 4 Preg-HC and 4 Preg-R samples for SC-RNA-seq. In total, we have obtained 45,859 high quality single cells for the cell clustering and gene expression analysis (Suppl. Fig. 2). Using ‘Seqgeq’ software, we first checked the library size versus genes expressed for high quality cells, secondly, we gated on high quality cells for total reads versus cells expressing genes and finally cells were gated for highly dispersed genes Suppl. Fig. 2b-c). Highly dispersed genes were used for PCA and PCA directed t-stochastic neighbour embedding (t-SNE) analysis – a statistical method visualizing high-dimension (Suppl. Fig. 2b-c). Further, using ‘Seurat’ plugin, we calculated the differential gene expression and cell clusters and explored different immune cell subsets (Suppl. Fig. 2d). We observed 20 cell clusters and have identified several major immune cells populations – monocytes, dendritic cells, CD4^+^ T cells, MAIT cells, CD8^+^ T cells, NK cells, B cells and megakaryocytes (Suppl. Fig. 3d). Additionally, we overlaid the Preg-HC or Preg-R samples on all the samples with different cell clusters. There were a few instances where cell clusters were less abundantly present like naïve CD4^+^ T cells, naïve B cells, NK cells and monocytes were reduced whereas MAIT cells were appeared to be present more in abundant in Preg-R samples compared with Preg-HC (Suppl. Fig. 2e). However, different subsets of CD4^+^ or CD8^+^ T cells were difficult to be distinguished.

Further, scRNA-seq analysis was performed using Seurat, performed the unsupervised clustering and visualized the data in Uniform Manifold Approximation and Projection (UMAP) plots. Cell clustering was based on PhonoGraph – a clustering algorithm, which is most robust when detecting refined subclusters (Liu *et al*, 2019). We discovered 34 cell subtypes of different immune cells (Fig. 4a). Further, individual known markers were defined to confirm the identity of a cell subtype group using unsupervised expression of each gene transcript (Fig. 4b-c). In detail, individual CD4^+^ T cells were clustered into naïve, central memory (CM), effector memory (EM), cytotoxic CD4^+^ T lymphocytes (CTL), SOX^+^ CD4^+^ T cells, and double negative (DN) cells. Similarly, CD8^+^ T cells were also classified in naïve, EM, CM, CTL and MAIT cells and NK cells into 5 different type of NK cell population based on distinct cell clusters (Suppl. Fig. 3a-c). Moreover, individual expression of total cell clusters for Preg-R and Preg-HC showed a significantly decreased number of specific cell subsets CD4^+^ CTL and B memory whilst CD8^+^ CTLs were significantly increased in numbers (Fig. 4d-g). In contrary, NK II and NK IV were nearly significantly changed, and NK II tended to be increased whilst NK IV tented to be decreased in Preg-R compared with Preg-HC (Fig. 4f).

**Fig. 4.**
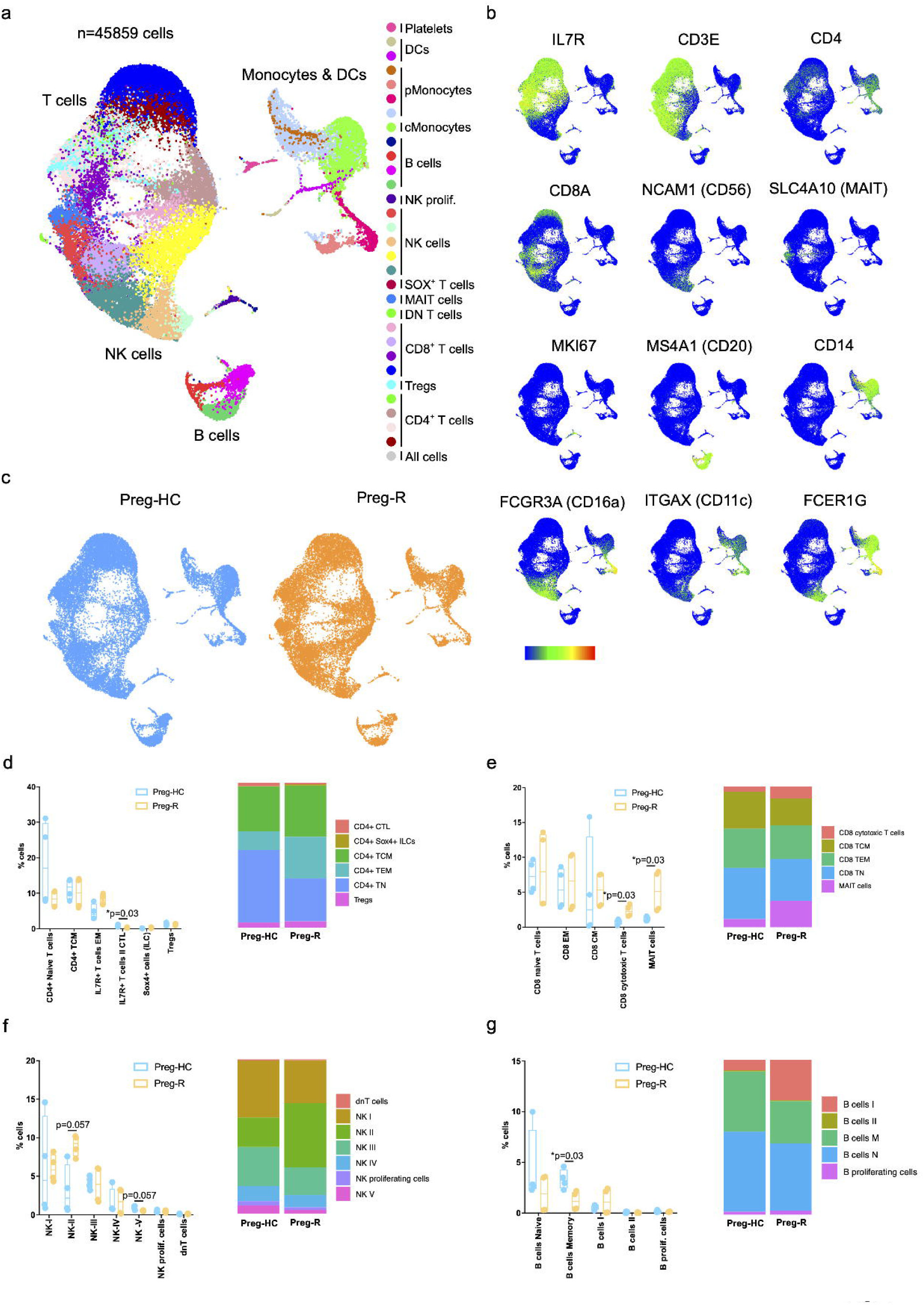
Dysregulated CD8, NK and B cells in recovered pregnant women a. SC-RNA-seq data were analyzed using Seqgeq software. Seurat and PhenoGraph pipelines were run to get the different subtype of immune cells. 13 different immune cells were identified based on distinct gene expression. b. Overlay of UMAP analysis of Preg-HC and Preg-R patients. c. Key markers for the validation of different immune cell clusters d. Percentage of major cell types in Preg-HC and Preg-R patients.

### Differential pathway regulation in monocytes, B cells and CD4^+^ TEM

Monocytes are the key innate immune cells in defence as well as ‘cytokine storm’ during COVID-19 pathology. We observed a clear reduced number of monocytes in Preg-R compared with Preg-HC. Thus, we explored the differential gene expression in Preg-R compared with Preg-HC, our data revealed that 329 genes were upregulated whilst 222 genes downregulated (Fold change ≥0.5 and q value ≤0.05) in Preg-R women (Fig. 5a). Based on upregulated genes, we performed Gene ontology pathway analysis using Metascape. We identified that Preg-R women had increased inflammatory gene signature which are mostly related with cytokine production, cell activation and regulation of defence response. Further, GO pathways suggested that TNF signalling pathways, catabolic process, lipid and orexin receptor pathways were also involved (Fig. 5b).

**Fig. 5.**
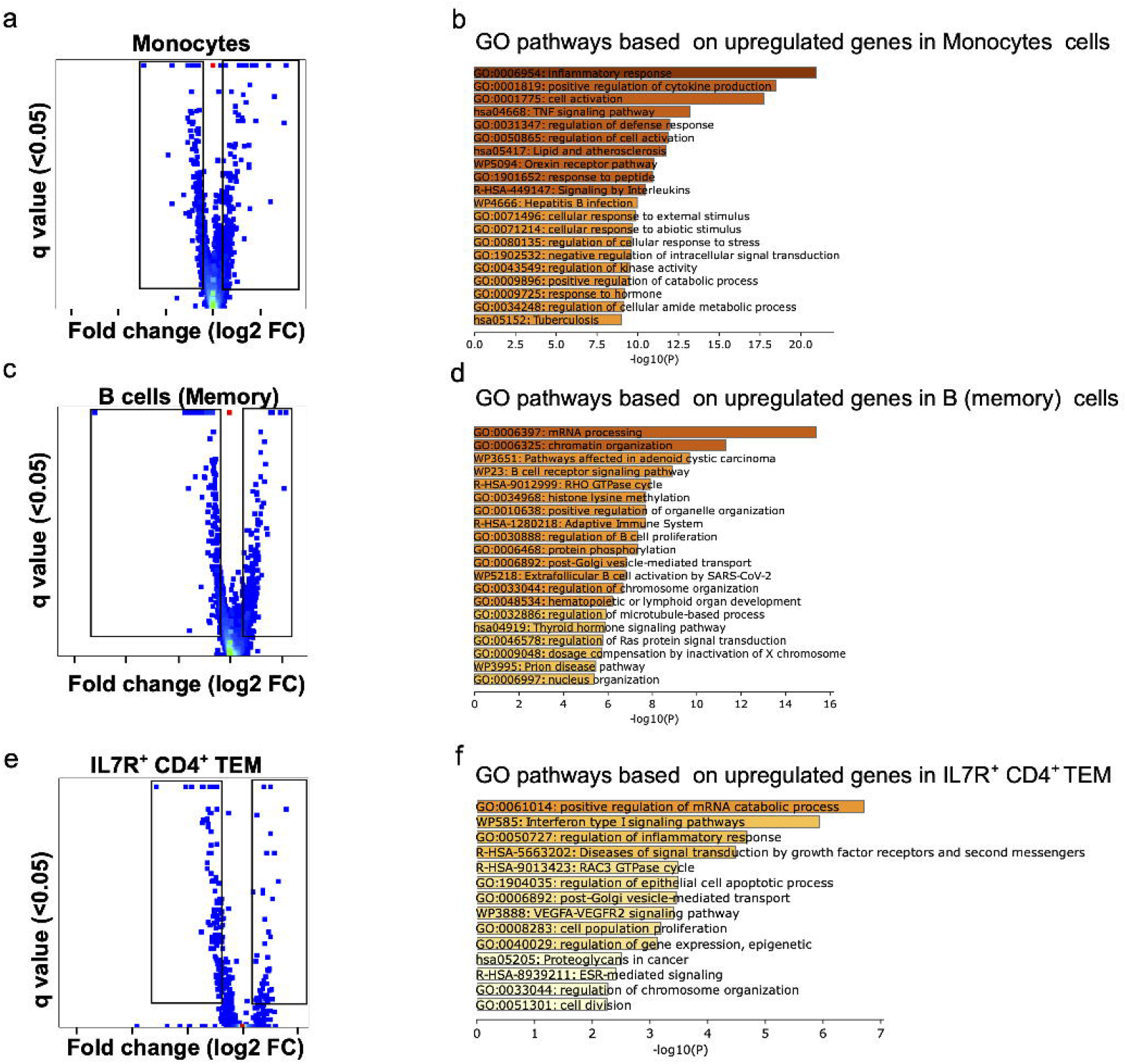
Inflammatory monocytes, activated memory B and CD4^+^ TEM cells in Preg-R a. Differential gene expression analysis in monocytes in Preg-R vs Preg-HC. Volcano plots showed the significantly upregulated and downregulated genes. b. GO pathway analysis based on upregulated genes in monocytes c. Gene expression analysis in B memory cells. d. GO pathways anaylsis based on upregulated genes. e. Differential gene expression analysis in IL-7R^+^ CD4^+^ TEM. Volcano plots represent significantly up and downregulation. f. GO pathways analysis based on upregulated genes in CD4^+^ TEM cells.

Further, B cells which are involved in humoral immune response were reduced in Preg-R and gene expression analysis highlighted that several genes were differentially regulated (349 upregulated and 229 downregulated) (Fig. 5c). The GO pathway analysis revealed that B cell receptor signalling pathways and extracellular B cell activation by SARS-CoV-2 were upregulated (Fig. 5d). Furthermore, GO pathways were related with mRNA processing, chromatin organization and histone lysine methylation which could be involved in imprinting of IgG antibody formation.

CD4^+^ TEM cells were also differentially abundant in Preg-R women, therefore, we also explored the differential (43 upregulated and 90 downregulated) gene expression and found that several genes were upregulated whilst 2-fold genes were downregulated (Fig. 5e). GO pathway analysis revealed that mRNA catabolic process was positively regulated, and further pathways related with regulation of inflammatory response, interferon type I signalling pathways, cell proliferation, apoptosis, cell division and epigenetic were upregulated (Fig. 5f).

### NK, MAIT and CD8^+^ T cell had reduced cytotoxic functions

Previously, we and others have shown that NK cell and CD8^+^ T cells were decreased in moderate and recovered COVID-19 patients respectively (Maucourant *et al*, 2020; Singh *et al*, 2021; Tian *et al*, 2021) and both cell types decreased with increased severity of the disease. MAIT cells were also involved in COVID-19 infection (Chen *et al*., 2021b). MAIT cells express the receptors for type I IFNs, IL-12, IL-15 and IL-18 thus these cells could potentially be activated by proinflammatory cytokines (Raffetseder *et al*, 2021). We therefore, explored the markers related with NK cells, MAIT and CD8^+^ CTLs functions in pregnant women recovered after SARS-CoV-2 viral infection. GO pathway analysis for NK type II cells based on upregulated genes (13 genes only) revealed upregulation of chromatin organization and regulation of cell adhesion molecules, whist based several down regulated gene were related with perforin and granzymes (Fig. 6a, b). GO enrichment pathways (based on 82 downregulated genes) revealed that several pathways - cell activation, cytokine signalling, natural killer cell mediated cytotoxicity, and oxidative phosphorylation, SARS-CoV-2 signalling pathways were downregulated. These pathways have some common genes such as GZMB, IL32 and IRF9 which were downregulated in Preg-R women compared with Preg-HC (Fig. 6c, d). In a similar fashion in NK type IV cells, mRNA metabolic pathway, cytokine signalling, cellular response to stress and hypoxia were upregulated (based on 264 genes) whilst, antigen processing and presentation, natural killer cell mediated cytotoxicity, neutrophil degranulation, SARS-CoV-2 network signalling pathway, interferon alpha/beta signalling and cytokine signalling were downregulated (218 genes) in Preg-R compared with Preg-HC (Fig. 6e-g). The most common genes related with cytotoxic functions and cell activation including CD2, FCFR3A, KLRC2, TUBB, IL2RB, AREG, GNLY, GZMK and GZMA were significantly downregulated in Preg-R compared with Preg-HC (Fig. 6h).

**Fig. 6.**
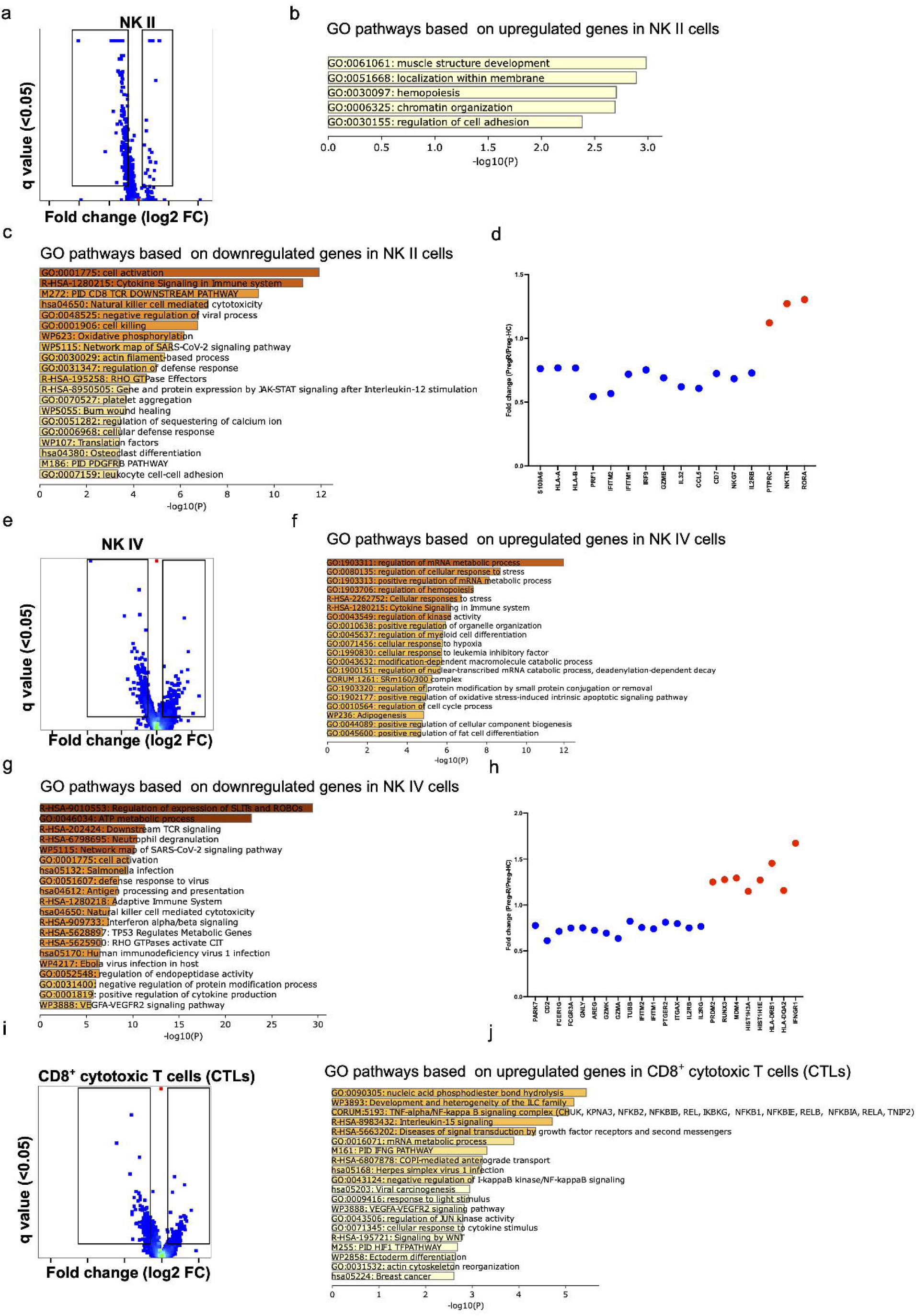
Reduced cytotoxic functions of NK and CD8^+^ CTLs in Preg-R a. Differential gene expression analysis in Preg-R of NK II cells. Most significantly genes are shown on the man. b. GO pathways based on upregulated genes in NK II. c. GO pathways based on downregulated genes in NK II. d. Dot plots represented significantly up and downregulated genes (selected genes) in NK II. e. Differential gene expression analysis in Preg-R of NK IV cells. Most significantly genes are shown on the map. f. GO pathways based on upregulated genes in NK IV. g. GO pathways based on downregulated genes in NK IV. h. Dot plots represented significantly up and downregulated genes (selected genes) in NK I. i. Differential gene expression analysis in Preg-R of NK IV cells. Most significantly genes are shown on the man. j. GO pathways based on upregulated genes in NK VI.

MAIT cells which are involved in direct recognition of peptide without MHC molecules were also differentially expressed genes in Preg-R (250 genes up and 162 genes downregulated) compared with Preg-HC (Suppl. Fig. 4a). Several GO pathways were upregulated in Preg-R compared with Preg-HC which are related with mRNA metabolic progress and cellular response to stress as well as cell adhesion. More interestingly, inflammatory, and anti-inflammation pathways such as IL-1, IFNs type I and TGB-b receptor signalling were also upregulated in Preg-R compared with Preg-HC (Suppl. Fig. 4b).

Moreover, in case of CD8^+^ CTLs cells, several cytokines signalling pathway related were upregulated whilst, cytotoxic function pathway genes were downregulated. GO pathway analysis revealed that inflammation related pathways including TNF/NFkB, IFNG, VEGFA-VEGFR2, HIF and actin cytoskeleton pathways were upregulated (Fig. 6i-j), whilst adaptive immune system, type II interferon signalling, antigen processing and presentation of exogenous peptide antigen via MHC I, neutrophil degranulation and leukocyte mediated cytotoxicity were downregulated in Preg-R. The most common gene related with cytotoxic functions CD2, GZMK, HLA-A, KLRK1, KLDR1, GZMK, GZMA, GZMM, IL-32 and IL2RG were significantly downregulated in Preg-R compared with Preg-HC (Suppl. Fig. 4c-d).

### Attenuated antiviral activity in recovered pregnant women

Previously, it was reported that after SARS-CoV-2 viral infection immunity is decreased in COVID-19 patients (Chua *et al*, 2020; Kim & Shin, 2021), thus, we investigated the status of their immune system. We discovered that expression of type I interferons (IFNs) receptors IFNAR1 and IFNAR2 was overall decreased in all immune cells, however IFNAR1 appeared to drastically decrease in B cell population (Fig. 7a). Downstream of type I IFNs signalling pathways TYK2, JAK1, STAT1, IRF9, ISG20, ISG20L2 and OAS3 were decrease in Preg-R patients (Fig. 7a). Type II IFNs signalling molecules such as IFN-g were increased in cytotoxic T cells. IFNGR1 and TNF were decreased in monocytes whereas cytotoxic CD8^+^ T cells and NK cells IFGR1 and TNF were increased in Preg-R patients compared with Preg-HC (Fig. 7b). JAK2 was also increased in monocytes in Preg-R patients compared with Preg-HC (Fig. 7b). Finally, TGFBI is upregulated in both CD8^+^ T and NK cells in Preg-R compared with Preg-HC (Fig. 7b). Overall, the data suggested a decrease of pro-inflammatory signalling mechanism and upregulation of anti-inflammatory molecules in Preg-R patients.

**Fig. 7.**
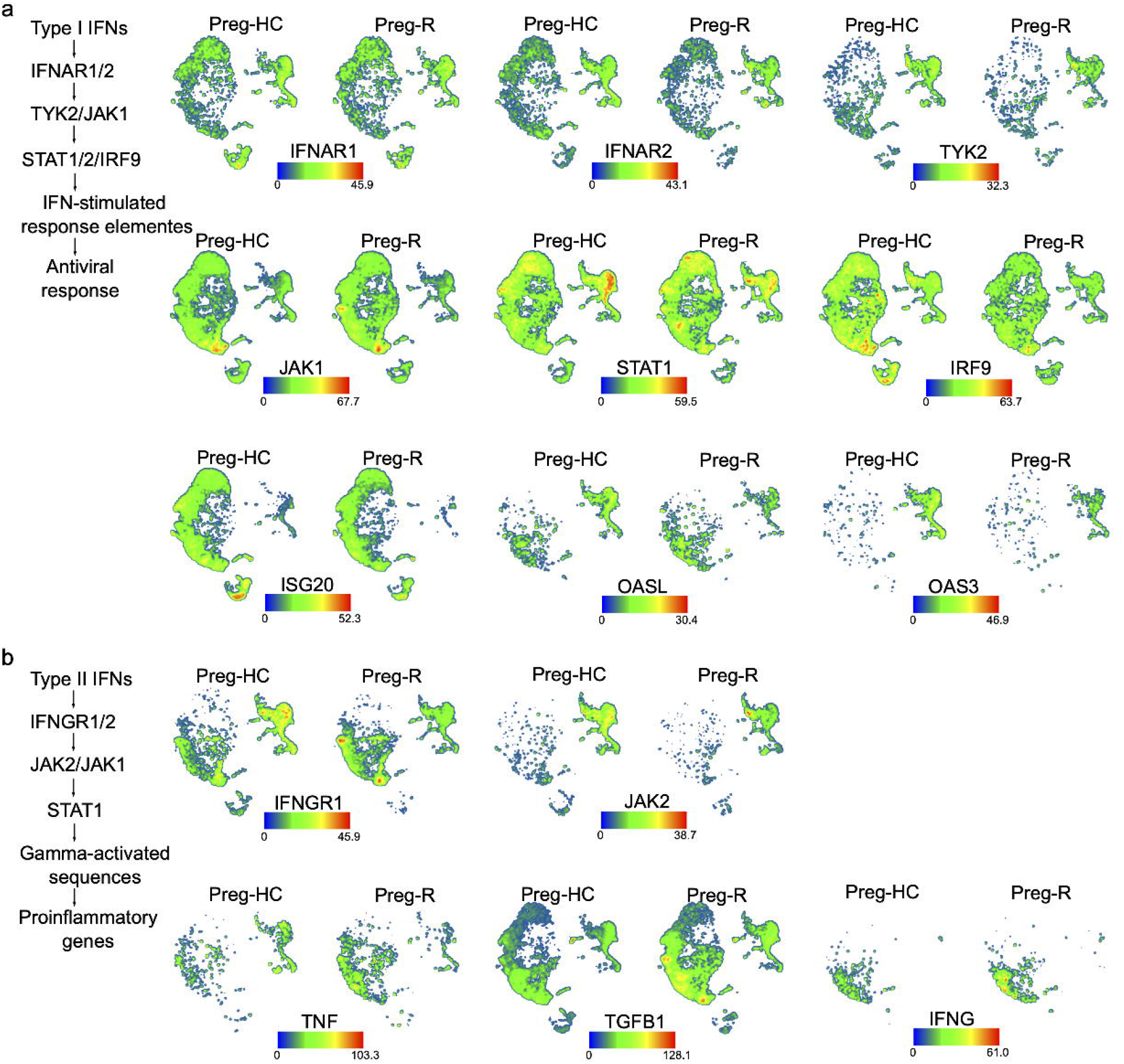
Dysregulated Type I and type II IFN signalling in recovered pregnant women. a. Type I IFN signalling genes (IFNAR1, IFAR2, TYK2, JAK1, STAT1, IRF-9, ISG20, ISG20L2, OAS3 and OASL) and their corresponding expression level. b. Type I IFN signalling gens (IFNG, IFNGR1, JAK2, TNF and TGFBI).

## Discussion

An immune-mediated cytokine response is the key driver for the severity of COVID-19 and resulting in a ‘cytokine storm’. In this report, using multi-colour flow cytometry and SC-RNA-seq, we have identified the immune signature during SARS-CoV-2 infection and in COVID-19 recovered pregnant women.Our multi-colour immunophenotyping data suggest that there was significant reduction in percentage of total lymphocytes whilst there was trend of decreased monocytes in SARS-CoV-2 infected and recovered pregnant women compared with healthy pregnant women. Our findings are further supported by recent data which also suggested that in pregnant recovered women there is a reduced proportion of lymphoid cells compared with healthy pregnant women (Chen *et al*., 2021b). Further, characterization of CD4^+^ T cells or CD8^+^ T cells revealed an increased percentage of early or late effector CD8^+^ T cells in pregnant SARS-CoV2 infected compared with pregnant recovered women. A similar trend was also observed for the CD4^+^ T cells effector cells. However, a reduced trend of effector cells either CD4^+^ or CD8^+^ T cells were also observed when comparisons were made between healthy control pregnant with SARS-CoV-2 pregnant women. These results imply that viral infection leads to transient reduction of effector cells which are increased in proportion once the viral is cleared. Our data agree with published findings which implied that moderate COVID-19 patients had increased reappearance of effector T cells compared with severe COVID-19 patients (Odak *et al*, 2020).

NK cells tended to be decreased in SARS-CoV2 and recovered pregnant compared with healthy pregnant women. Earlier studies reported no significant change in NK cells and suggested increased trend of NK cells in pregnant COVID-19 women (Chen *et al*., 2021a). Overall, our phenotyping data suggest decreased effector T cells and NK cells could stop an exacerbated immune response during the active infection of pregnant COVID-19 women. Cytokine and chemokines detection in the plasma of pregnant SARS-CoV-2 infected women revealed reduced levels of IL-12(p70) and increased RANTES levels (Chen *et al*., 2021a). These findings suggest that suppressed cytokine and chemokines levels could be helpful to avoid adverse outcome and have unique anti-SARS-CoV-2 response during pregnancy.

Furthermore, scRNA-seq revealed increased levels of effector CD4^+^ or CD8^+^ T cells in recovered pregnant compared with healthy pregnant women. Differential gene expression analysis of CD4^+^ TEM in recovered pregnant women suggested that several genes with interferon type I and inflammatory signalling pathways were changed. It appears that effector CD4^+^ T cells from recovered pregnant women has a strong protective effective arm, which may lead to a competitive advantage to avoid overt inflammation to protect the ongoing pregnancy. Further, CD4^+^ cytotoxic T cells were decreased whilst CD8^+^ cytotoxic T cells were significantly increased. Differential gene expression analysis of CD8^+^ CTL suggested that several pathways related with cytokine signalling (TNF, IFNG and IL-15), VEGFA-VEGFR2, actin cytoskeleton and HIF1 pathways were upregulated in recovered pregnant women. Furthermore, MAIT cells also have increased mRNA metabolic, IL-1, chromatin organization, interferon type I, TGFB, histone modification and immune system development were upregulated. pMonocytes also had increased inflammatory response, positive regulation of cytokine and cell activation and regulation of defence response. In a similar fashion, in B memory cells from recovered pregnant women also had increased chromatin organization, histone lysine methylation and activation of extrafollicular by SARS-CoV-2 and regulation of B cell proliferation were upregulated. Thus, scRNA-seq analysis suggested that a clear molecular advantage of infected recovered women to avoid the pregnancy complications. However, further large-scale studies are warranted to confirm these data and findings.

### Limitation of the study

Our results may have provided key information for future research work into how pregnant women are affected by SARS-CoV-2 infection. However, there are still some limitations to this study. First, our cohort sample size was relatively small as few pregnant women agreed to donate the blood samples to the transcriptomics study. Additionally, we were not able to recruit pregnant women with severe symptoms. Also, pregnancy state has a dynamic and evolving immune changes at different trimester ages, which could also affect the immune response to COVID-19 as the pregnant patients who enrolled in our research study mostly developed symptoms in the second and third trimester rather than in the first trimester.

## Figure legends

Sup. Fig.1 Characterization of PBMCs from pregnant infected and recovered from SARS-CoV-2 infection

a. Gating strategy for the 14-colour flow cytometry panel. Based on FSC and SSC we removed the cell debris and gated on live cells. FSC-A and FSC-W was used to remove the doublets and focussed on single cells. In the next step, we removed the cells using Violet Live/dead staining. Finally, we again we used FSC-A vs SSC-A to gate lymphocytes and monocytes based on size and granularity. Lymphocytes were discriminated in CD19 and CD3 based on cell surface markers and CD3 cells were discriminated into CD4 and CD8 cells using CD4 and CD8a antibodies.

b. FSC-A vs SSC-A show the monocytes and lymphocyte gated population (upper side FACS plots). The percentage of lymphocytes and monocytes shown by violin plots (lower side) for Preg-HC, Preg-SARS-CoV-2 and Preg-R samples.

c. FACS plots for CD4 versus CD8a staining gated on CD3+ T cells (upper side). The percentage of CD4^+^ and CD8a^+^ T cells in Preg-HC, Preg-SARS-CoV-2 and Preg-R samples.

Suppl. Fig. 2 Quality check and t-SNE analysis for single cell RNA-seq data from healthy control and recovered pregnant women

a. Total 56,586 cells were recovered after sequencing from 8 samples (Preg-HC and Preg-R). First RNA-sequenced cells were gated for genes expressed vs library size, we found 45,859 high quality cells. Next high QC cells were analysed for cells expressing based on total read cut off min cells 5 were chosen for further analysis. Cells expressing highly dispersed gene were gated and used for PCA and t-SNE analyses.

b. Highly dispersed genes were subjected to PCA analysis, and we ran 25 principle components (PC).

c. PC guided unsupervised clustering analysis was used calculating t-SNE map.

d. For identification of cell clusters from t-SNE plots, we identified 3 major clusters – monocytes, lymphocytes and B cells as shown in arbitrary clusters. Further each cluster based on RNA transcript levels and divided into major cell types – CD4+ T cells, CD8+ T cells, NK cells, NKT, ILC/MAIT, p and cMonocytes, dendritic cells and megakaryocytes.

e. Overlay of pregnant healthy control and recovered pregnant t-SNE clustering show difference in cell clustering.

Suppl. Fig. 3 Representation of individual clusters based on phonograph-based cell clustering

a. CD4^+^ T cell cluster. 7 major subtypes of CD4^+^ T cells were identified.

b. CD8^+^ T cell cluster. 5 subtypes of CD8^+^ T cells were defined based on gene expression.

c. Five subtypes of NK cells in pregnant healthy and COVID-19 recovered patients.

Suppl. Fig. 4 Deregulated MAIT and CD8^+^ CLTs response in Preg-R

a. Differential gene expression analysis in Preg-R of MAIT cells. Most significantly genes are shown on the volcano plot.

b. GO pathways based on upregulated genes in MAIT cells.

c. GO pathways based on downregulated genes in CD8^+^ CTLs cells.

Dot plots represented significantly up and downregulated genes (selected genes) in CD8^+^ CTLs (Red = upregulated, Blue = Downregulated (0.005)).

## Material and methods

### Ethics statement for the study participants

This study is part of the overall study of the transcriptomic and protein analysis of pregnant women with a history of COVID-19 infection at the epicentre of the COVID-19 pandemic in Malaysia approved by Research Ethics Committee, National University of Malaysia (JEP-2021-465) and from the Medical Research Ethics Committee (MREC) of Ministry of Health Malaysia (ID-58736). This was a cross-sectional study carried out in Malaysia with a total of 21 pregnant women fulfilling the inclusion and exclusion criteria were selected using convenient sampling. 4 pregnant women infected with SARS-CoV-2 virus, 4 pregnant women who recovered from COVID-19 and 13 pregnant women from healthy controls were provided written informed consent to participate in this study. Disease severity for this study was determined by symptoms. These women were recruited from the obstetrics and gynaecology clinic, outpatient clinical wards from April 2020 until Feburary 2021. The study procedures were carried out in accordance with Declaration of Helsinki.

### Cohort size and sample collection

We collected blood from total 8 SARS-CoV-2 infected and recovered pregnant women (n=4/group) and 13 pregnant women enrolled in the Department of Obstetrics and Gynaecology, Universiti Kebangsaan Malaysia Medical Centre, Kuala Lumpur, Malaysia. Written informed consent from patients was obtained (Universiti Kebangsan Malaysia). Eligibility criteria included age >18 years and positive RT-PCR test for infected and negative RT-PCR test for healthy control and recovered pregnant women. To protect the identity of the enrolled pregnant women, pseudonymized samples were sent to Tübingen University for single cell RNA-sequencing (scRNA-seq) processing and analysis. Blood was collected from pregnant women infected with SARS-CoV-2 virus 1-3 days after their hospital admittance or prior their caesarean section delivery. From pregnant women recovered from COVID-19 who were admitted in wards before delivery and pregnant women from healthy controls who attended the outpatient clinic, 3-5 ml blood was collected in a 5 ml plain and a 10 ml Lithium heparin vaccutainer tubes. Serum isolation via centrifugation will be conducted immediately after collection. Human Peripheral Blood Mononuclear Cell (PBMCs) were isolated by the standard Ficoll method (Singh *et al*., 2021).

### Preparation of PBMCs for single cell RNA sequencing (scRNA-seq) and antibody (surface and intracellular proteins) staining for flow cytometry

Frozen PBMCs were thawed at 37 ^0^C in a water bath for 2 minutes or until a small ice crystal remains. In a biosafety hood (Level 2), thawed cells slowly transferred to a 50 ml conical tube using a wide-bore pipette tip and rinsed the cryovial with 1 ml warm complete RPMI1640 medium (RPMI1640, 10% FBS, anti/anti) dropwise (1 drop per 5 seconds) to the 50 ml of tubes while gently shaking the tube. Further, sequentially diluted cells in the 50 ml tube by incremental 1:1 volume addition of complete RPMI1640 medium for a total of 5 times with 1 minute wait between each addition (total volume 32 ml). After completion of adding the medium, tube was centrifuged at 400xg for 5 minutes at room temperature. Supernatant was discarded after centrifuge leaving 1 ml of medium was left and mixed the cells using regular bore pipette or 10ml of Pasteur pipette tip. After mixing, 19ml of complete RPMI1640 was added and washed the cells at 400xg for 5 minutes at room temperature to remove any remaining DMSO from the cells. Again, supernatant was discarded, and cells were resuspended in 2 ml of medium and cells were counted. After counting the cells 1×10^6^ cells were used for each 14 colour FACS panel and 0.5×10^6^ cells were used for single-cell RNA sequencing (SC-RNA-Seq).

### Flowcytometry data analysis staining and data analysis

For flow cytometry staining, 1×10^6^ cells were taken in 96 well plate and washed with DPBS (Ca^2+/^Mg^2+^ free). First, the cells were stained with 5 ul of live and dead dye (1:400 dilution in PBS) and stained the cells for 15 minutes in dark at room temperature. After incubation, cells were washed once with PBS and added 50ul of PBS into each well. 12 colour antibodies cocktail was made for the surface recognizing markers (CD3, CD4, CD8, CD19, CD56, HLA-DR, CD38, CD154, CD45R, CCR7, CD14, CD16) as reported earlier (Singh *et al*., 2021). For each reaction, we used 2.5ul antibody as well as 5.0ul of superbright staining buffer (to distinguish two different antibody in the superbright colour). Cells were incubated with antibody cocktail for 30-40 minutes. After incubation, cells were washed with PBS and fixed using fix/permb buffer for 45 minutes. After fixation, cells were permeabilized using 1x permeabilization buffer and added 2ul of Foxp3 intracellular antibody and incubated at room temperature for 30-40 minutes. After incubation, cells were washed with PBS and cell samples were acquired on BD Fortessa flow cytometry. Data were analysed using Flow jo for 2D FACS plots dimensional reduction methods.

### Sample preparation of SC-RNA-seq

Thawed 0.5×10^6^ cells washed 3x at 400xg for 5 minutes at room temperature with 1x staining buffer (Biolegend) and kept on ice during the live dead staining using acridine orange and propidium dye to measure the cell viability before proceeding for 10x experiments. We followed the 10x chromium SC-RNA-seq protocol. In brief, after measuring the cell viability, cells were loaded on 10x Chromium Chip.

Single cells were prepared in the Chromium Single Cell Gene Expression Solution using the Chromium Single Cell 3’ Gel Bead, Chip, and Library Kits v1 (10x Genomics) as per the manufacturer’s protocol. In all, 20,000 total cells were loaded to each channel with an average expected recovery of 6000-9000 cells. The cells were then partitioned into Gel Beads in Emulsion in the Chromium instrument, where cell lysis and barcoded reverse transcription of mRNA occurred, followed by amplification, shearing, and 3’ adapter and sample index attachment. Libraries were quantified by QubitTM 2.0 Fluorometer (ThermoFisher) and fragment size was controlled using 2100 Bioanalyzer with High Sensitivity DNA kit (Agilent). Sequencing was performed in paired-end mode with a S1 and S2 flow cell (100 cycles) using NovaSeq 6000 sequencer (Illumina) at NCCT, IMGAG, Tübingen, Germany.

For the 10× Genomics sequencing data alignment and quantification, the sequencing data were processed using the CellRanger software (v3.0.1) with default parameters and the GRCh38 v3.0.0 human reference genome.

### scRNA-seq analysis by Seqgeq software (BD Bioscience) using Seurat and associated plugin

First, cell ranger filtered feature matrix files (.HD5 format) were concatenated in “Seqgeq” online browser user friendly software (BD Bioscience and Flow Jo). We used n=4 Preg-HC and n=4 Preg-R (in duplicate) samples for analysis (total cells recovered from sequencing; n=55,588). Before concatenations each matrix file was normalized (counts per 10,000) and adjusted transformation to reflect normalized ranges. Concatenated file was normalized (counts per 10,000) and performed the quality check (QC) as described online (https://www.flowjo.com/learn/flowjo-university/seqgeq). In brief, we firstly identified the library size and gene expressed in total to decide the good quality cells in cell view platform. We identified 81% cells (n=45,859) were good quality (Suppl. Fig 2) and use for deciding the gene cut off for the upper and lower limit and gated as high parameters in cell view platform. High quality gated genes were selected for highly dispersed genes. These dispersed genes (n=1200) were used for software inbuilt dimensional reduction platforms - principal component analysis (PCA) guided t-distributed stochastic neighbour embedding (t-SNE) analysis to present the data into biaxial plots, while preserving variance contained in biological populations. Further, “Seurat plugin” analyses to perform the unsupervised cell clustering and UMAP analysis including the if any batch corrections. The Seurat output resulted in 20 cell clusters. Cell clusters were further defined on Seurat UMAP using another plugin “Phenograph” which results in 31 cell clusters. Each cell cluster was defined on differential gene expression analysis with respected to the all the cell clusters.

### Psuedotime trajectory of B cells

Monocle plugin was used for pseudotime trajectory to understand the B cell development. B cells were concatenated from Seurat output pipeline and were fed into Monocle plugin seqgeq software as default setting.

### Differential gene expression in Preg-HC vs Preg-R

Each cell clusters were compared for differential gene expression Preg-R vs Preg-HC to identify the upregulated and downregulated gene expression for gene enrichment analysis. Volcano plots were used for identifying differentially regulated genes to asecertain genes expressed abov a certain fold change and statistically significant p-value. By default, p-Values are adjusted using False Discovery Rate (FDR) is calculated according to the Benjamini-Hochberg algorithm and considered significant (q value ≤0.05).

### Gene ontology (GO) pathway enrichments using Metascape

Differentially upregulated or downregulated genes from each cell clusters were used for GO pathway enrichment anlysis using online Metascape tool (https://metascape.org). In brief, Metascape first automatically converts the input identifiers (Entrez Gene ID, RefSeq, Ensembl ID, UnProt ID or Symbol) into Human Entrez Gene ID. We first identified all statistically enriched terms (can be GO/KEGG terms, canonical pathways, hall mark gene sets, etc., based on the default choices under Express Analysis). Accumulative hypergeometric p-values and enrichment factors were calculated and used for filtering and the remaining significant terms were then hierarchically clustered into a tree based on Kappa-statistical similarities among their gene memberships (used in NCI DAVID site). Then 0.3 kappa score was applied as the threshold to cast the tree into term clusters. The terms within each cluster are exported in the Excel spreadsheet named “Enrichment Analysis” and presented in bar diagram for selected pathways.

### Statistics analysis

Cytometry data was analyzed using FlowJo 10.8.1. Statistical analyses were performed in GraphPad Prism 9.3.0, unless otherwise stated. The statistical details of the experiments are provided in the respective figure legends. Data plotted in linear scale were expressed as Mean ± Standard Deviation (SD). Mann–Whitney U or Wilcoxon rank-sum tests with Dunn’s post-hoc (for multiple comparisons) were applied for unpaired comparisons, respectively.

## Author’s contributions

NHAA: Conceiving the study, blood sample collection, metadata collection, performing the experiments, data analysis

MSS: Conceiving the study, performing initial part of scRNA-seq sample preparation, data analysis, figure preparation, writing the manuscript

AKL: Single cell data analysis, discussion, preparation of the figures

MNS: Conceiving the study, metadata collection, funding acquisition

NMK, AK, and NAF: Blood sample collection, PBMCs preparation, metadata collection

SPS, OK: PBMCs preparation for the flow staining and acquisition of the samples on flow cytometry

EK: scRNA-seq data processing and data analysis

SO, NC, SYB, OR: Funding acquisition, provided tools for sc-RNA-seq, data analysis and discussion of the scRNA-seq data

DeCOI: tools and data discussion

YS: Conceiving the study, provide support for the experiments, data analysis, funding acquisition, writing the manuscript and overall project management

## Acknowledgements

We thank all the pregnant women who participated in this study. All the lab members who participated in fruitful discussion and technical assistance.

## DeCOI consortium

DeCOI members are presented in https://decoi.eu/members-of-decoi/

## Funding for the study

This project is supported by Ferring Pharmaceuticals to YS. MSS and MNS were also independently supported by Ferring Pharmaceuticals to provide materials and tools for this project. NGS sequencing methods were funded and performed with the support of the DFG-funded NGS Competence Center Tübingen (INST 37/1049-1) and Project-ID 286/2020B01 – 428994620. MSS is supported by the Margarete von Wrangell (MvW 31-7635.41/118/3) habilitation scholarship co-funded by the Ministry of Science, Research, and the Arts (MWK) of the state of Baden-Württemberg and by the European Social Funds.

## Conflict of Interest

The authors declare that the research was conducted in the absence of any commercial or financial relationships that could be construed as a potential conflict of interest.

## Data and code availability

- The raw data generated by SC-RNA-sequencing from this study are available to download through the public repository via the following accession number or link:
- This paper does not report any original code.
- Any additional information required to re-analyze the data reported in this paper is available from the lead contact upon reasonable request.

## Lead contact

Requests for resources and reagents and for further information should be directed to and will be fulfilled by the Lead contact

## Notes

### Competing Interest Statement

The authors have declared no competing interest.

