## Supplementary material for "Single-cell RNA sequencing highlights a reduced function of natural killer and cytotoxic T cell in recovered COVID-19 pregnant women": Suppl. Info

^7^DeCOI members are presented in <https://decoi.eu/members-of-decoi/>

^#^Equal contributions

*Lead Contact for the study

Dr Yogesh Singh

Institute of Medical Genetics and Applied Genomics

NGS Competence Centre Tübingen (NCCT)

Research Institute of Women’s Hospital

University of Tübingen

Calwerstraße 7, 72076, Tübingen

Germany

Key words: Pregnancy, COVID-19, PBMCs, NK cells and cytotoxic cells

Running title: Immune response in COVID-19 pregnant women

**Suppl. Figure legends**

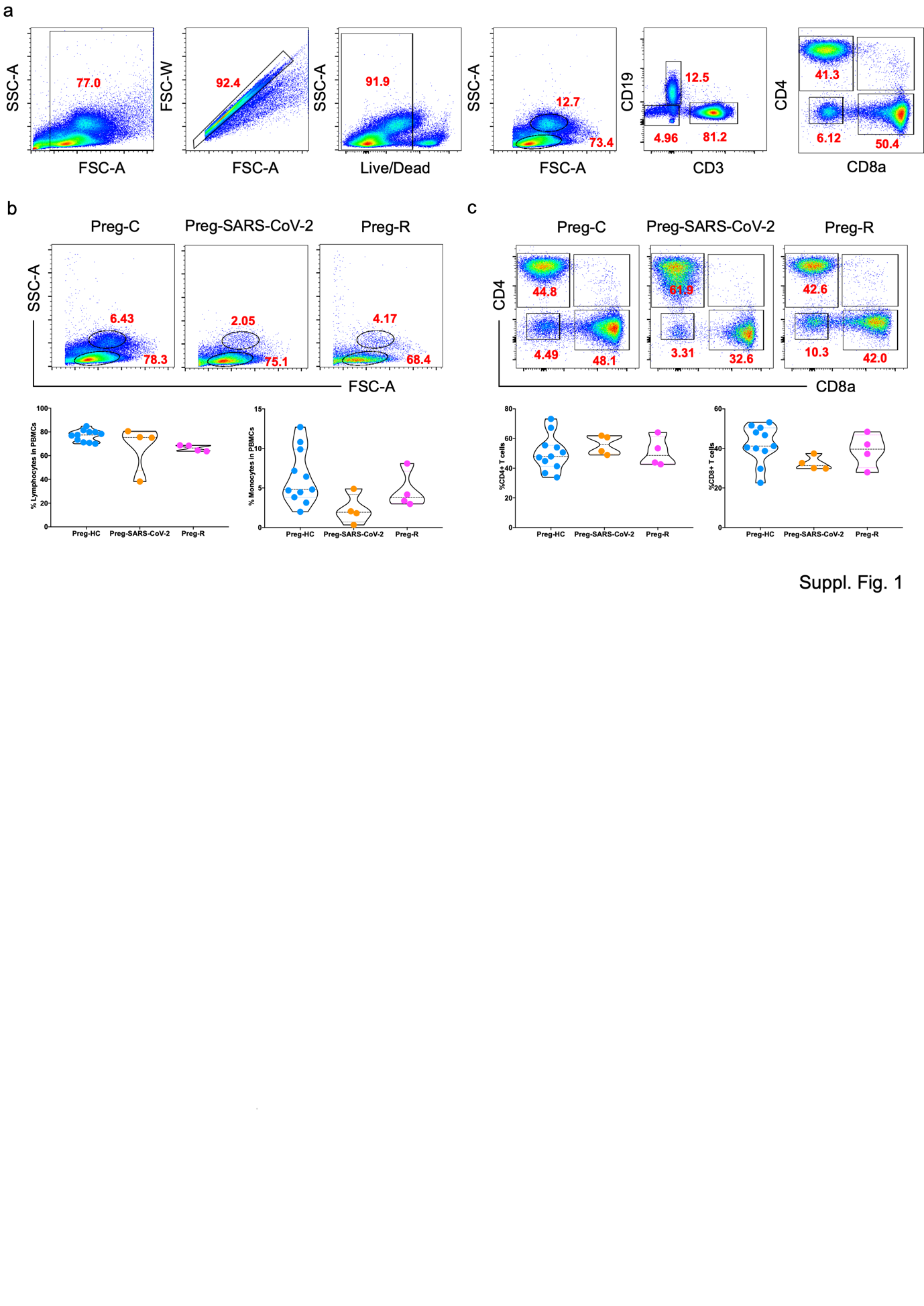

Sup. Fig.1 Characterization of PBMCs from pregnant infected and recovered from SARS-CoV-2 infection

a. Gating strategy for the 14-colour flow cytometry panel. Based on FSC and SSC we removed the cell debris and gated on live cells. FSC-A and FSC-W was used to remove the doublets and focussed on single cells. In the next step, we removed the cells using Violet Live/dead staining. Finally, we again we used FSC-A vs SSC-A to gate lymphocytes and monocytes based on size and granularity. Lymphocytes were discriminated in CD19 and CD3 based on cell surface markers and CD3 cells were discriminated into CD4 and CD8 cells using CD4 and CD8a antibodies.
b. FSC-A vs SSC-A show the monocytes and lymphocyte gated population (upper side FACS plots). The percentage of lymphocytes and monocytes shown by violin plots (lower side) for Preg-HC, Preg-SARS-CoV-2 and Preg-R samples.
c. FACS plots for CD4 versus CD8a staining gated on CD3+ T cells (upper side). The percentage of CD4^+^ and CD8a^+^ T cells in Preg-HC, Preg-SARS-CoV-2 and Preg-R samples.

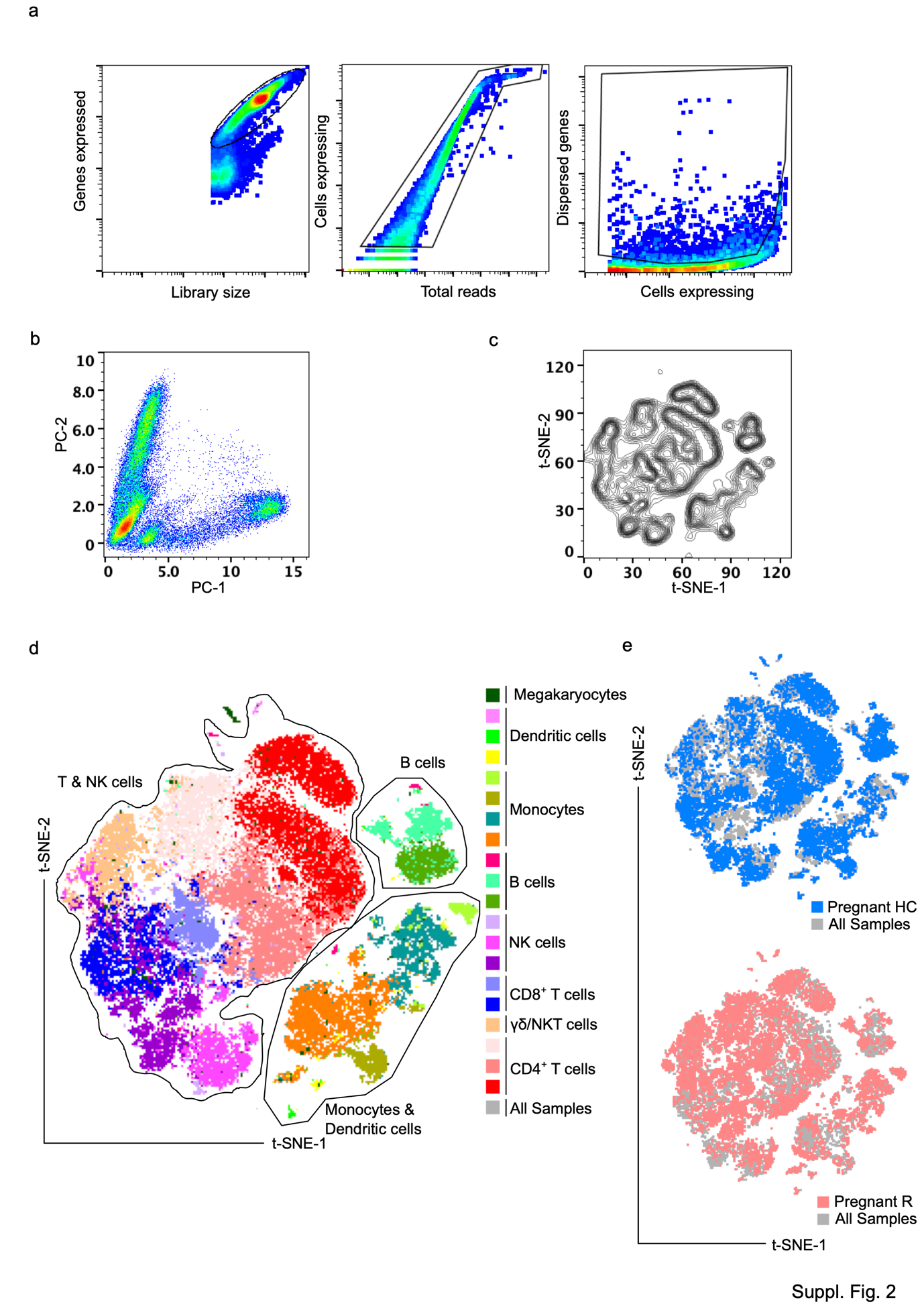

Suppl. Fig. 2 Quality check and t-SNE analysis for single cell RNA-seq data from healthy control and recovered pregnant women

a. Total 56,586 cells were recovered after sequencing from 8 samples (Preg-HC and Preg-R). First RNA-sequenced cells were gated for genes expressed vs library size, we found 45,859 high quality cells. Next high QC cells were analysed for cells expressing based on total read cut off min cells 5 were chosen for further analysis. Cells expressing highly dispersed gene were gated and used for PCA and t-SNE analyses.
b. Highly dispersed genes were subjected to PCA analysis, and we ran 25 principle components (PC).
c. PC guided unsupervised clustering analysis was used calculating t-SNE map.
d. For identification of cell clusters from t-SNE plots, we identified 3 major clusters – monocytes, lymphocytes and B cells as shown in arbitrary clusters. Further each cluster based on RNA transcript levels and divided into major cell types – CD4+ T cells, CD8+ T cells, NK cells, NKT, ILC/MAIT, p and cMonocytes, dendritic cells and megakaryocytes.
e. Overlay of pregnant healthy control and recovered pregnant t-SNE clustering show difference in cell clustering.

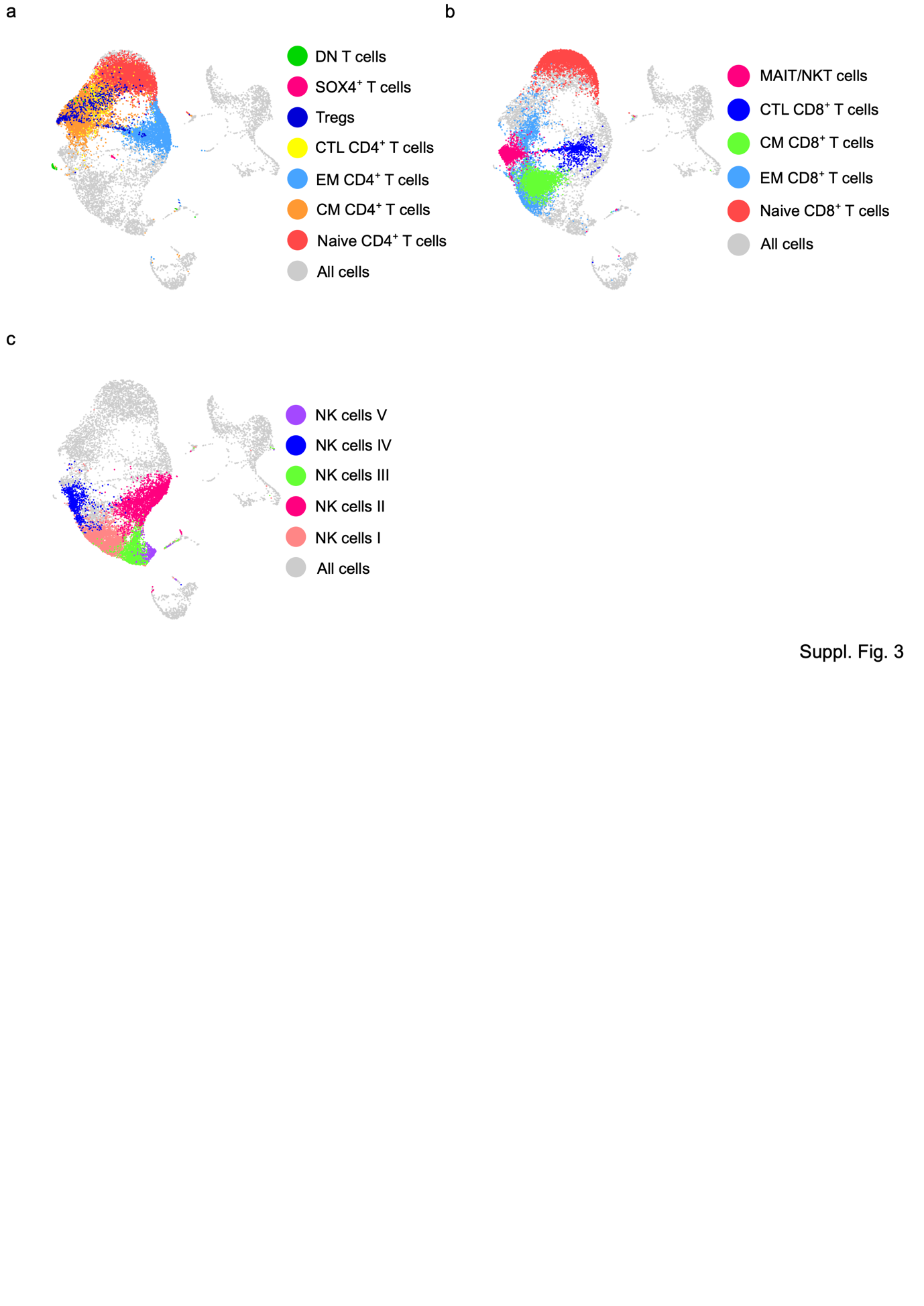

Suppl. Fig. 3 Representation of individual clusters based on phonograph-based cell clustering

a. CD4^+^  T cell cluster. 7 major subtypes of CD4^+^ T cells were identified.
b. CD8^+^ T cell cluster. 5 subtypes of CD8^+^ T cells were defined based on gene expression.
c. Five subtypes of NK cells in pregnant healthy and COVID-19 recovered patients.

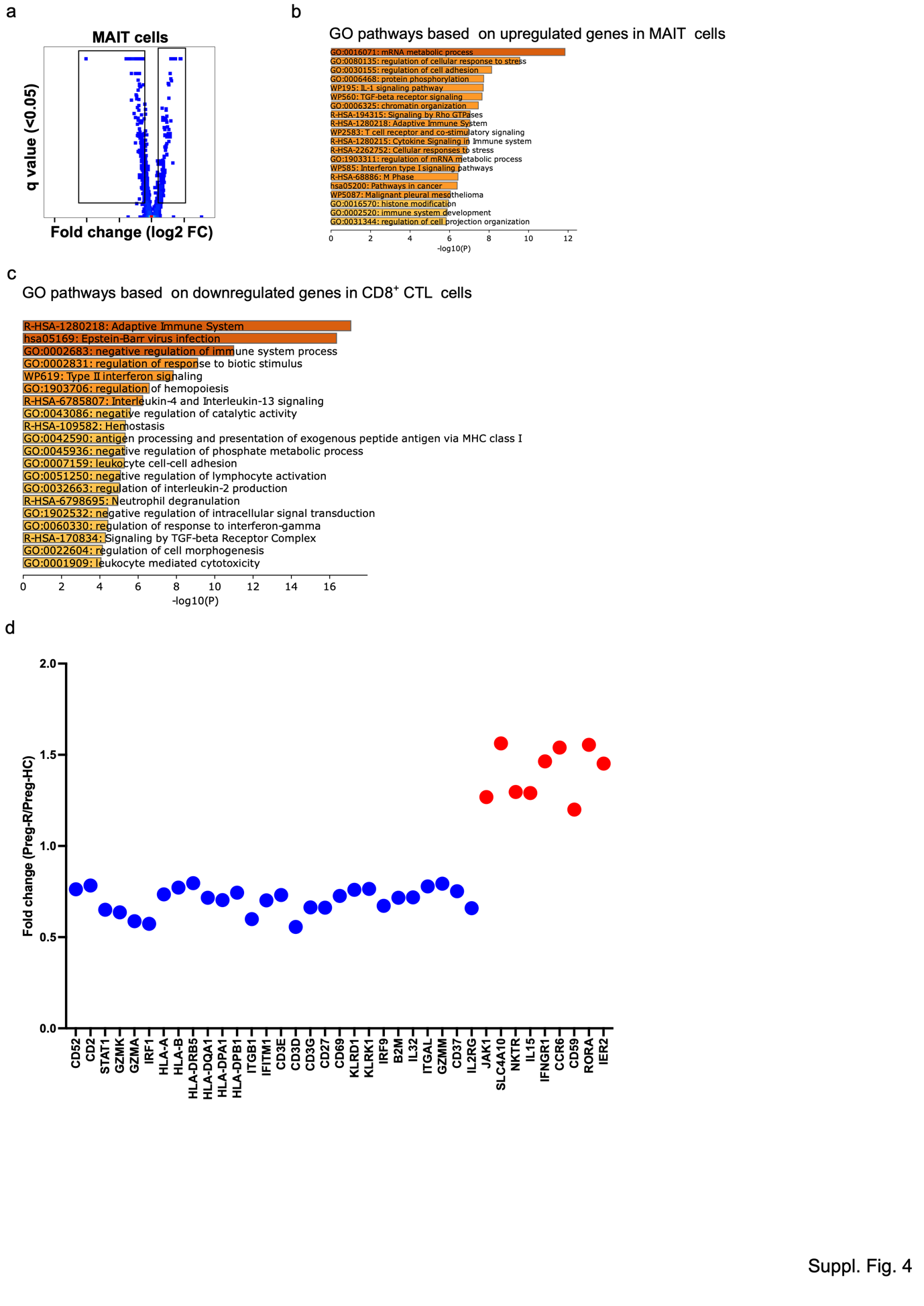

Suppl. Fig. 4 Deregulated MAIT and CD8^+^ CLTs response in Preg-R

a. Differential gene expression analysis in Preg-R of MAIT cells. Most significantly genes are shown on the volcano plot.
b. GO pathways based on upregulated genes in MAIT cells.
c. GO pathways based on downregulated genes in CD8^+^ CTLs cells.

Dot plots represented significantly up and downregulated genes (selected genes) in CD8^+^ CTLs (Red = upregulated, Blue = Downregulated ( 0.005)).
